# A paper-filter system to investigate the real micro-environment circuiting plant roots

**DOI:** 10.1101/553479

## Abstract

The micro-environment circling the plant root is an interesting topic for many researchers. Until now there is not an approach to investigate the exact density of chemicals surrounding the plant roots. here we use a simple paper filter system and gas chromatography to quantify the exact density of chemicals surrounding the plant roots. this can help solve a long-existing doubt about the ‘real micro-environment’ surrounding the plant roots.

---

There is a long history since the research about root executes in plants started in the end of 18^th^ century. In 1795, Plenk et.al found that the *Orobanche cumana Wallr* can only geminate with host roots, which led him to imagine that there are some executes from plant roots^1^. Since then there are huge number of papers about plant root executes^2-4^. In these papers many researchers found that plant root executes can have variable functions to plant germination and development. Owing to technical problem most of the time we can only detect the root executes in soil or do research with liquid culture media. Obviously the surface of the plant root is an executes layer which is a reservoir for root executes and sheds them to the liquid media or soil (figure1). There always exists a doubt: how much is the real density of the root executes circuiting the root? For the sake of this technical obstacle many researchers faced a big problem: they cannot decide the execute concentration in their liquid media when they want to exploit the influence of executes to plants because nobody knows the exact density circuiting the plant roots, which makes their results face many challenges. Here we want to introduce a simple method to measure the density of the thin layer circuiting the plant roots.

**Figure 1:**
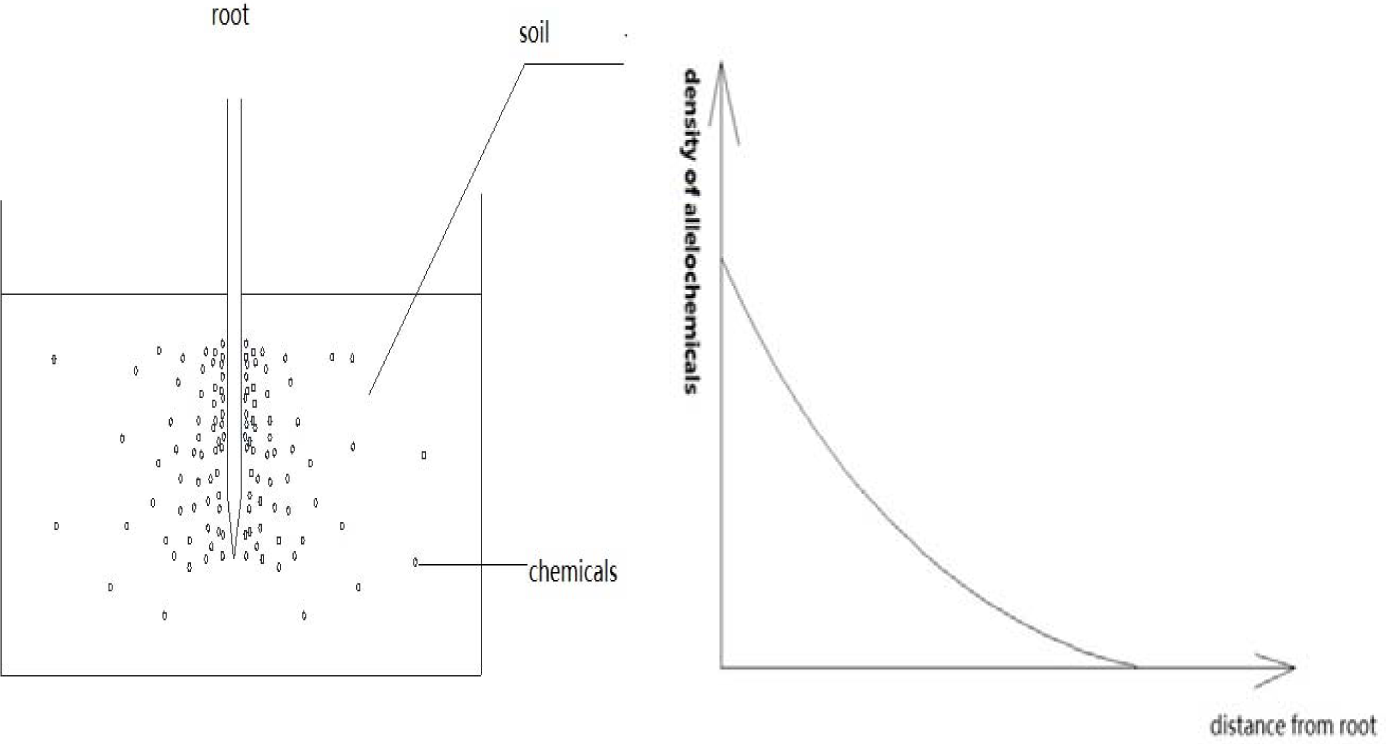
distribution of allelochemical in soil surrounding the root

## Construction of membrane-root system

At first we prepare 2 pieces of filter paper, which should be from the same company, with the same product lot number. The plants are cultured in water pot. When it arrives at expected period(the researchers will decide it according to their aim), we will collect it from the water pot and wrap one piece of filter paper circuiting the root tightly. A cotton thread may be necessary in this step. Then the plant will be replanted in soil, in the meanwhile another filter paper is also put into the soil, distance from the plant is 8-10 cm. The soil should be the object soil.(fig.2)

**Figure 2:**
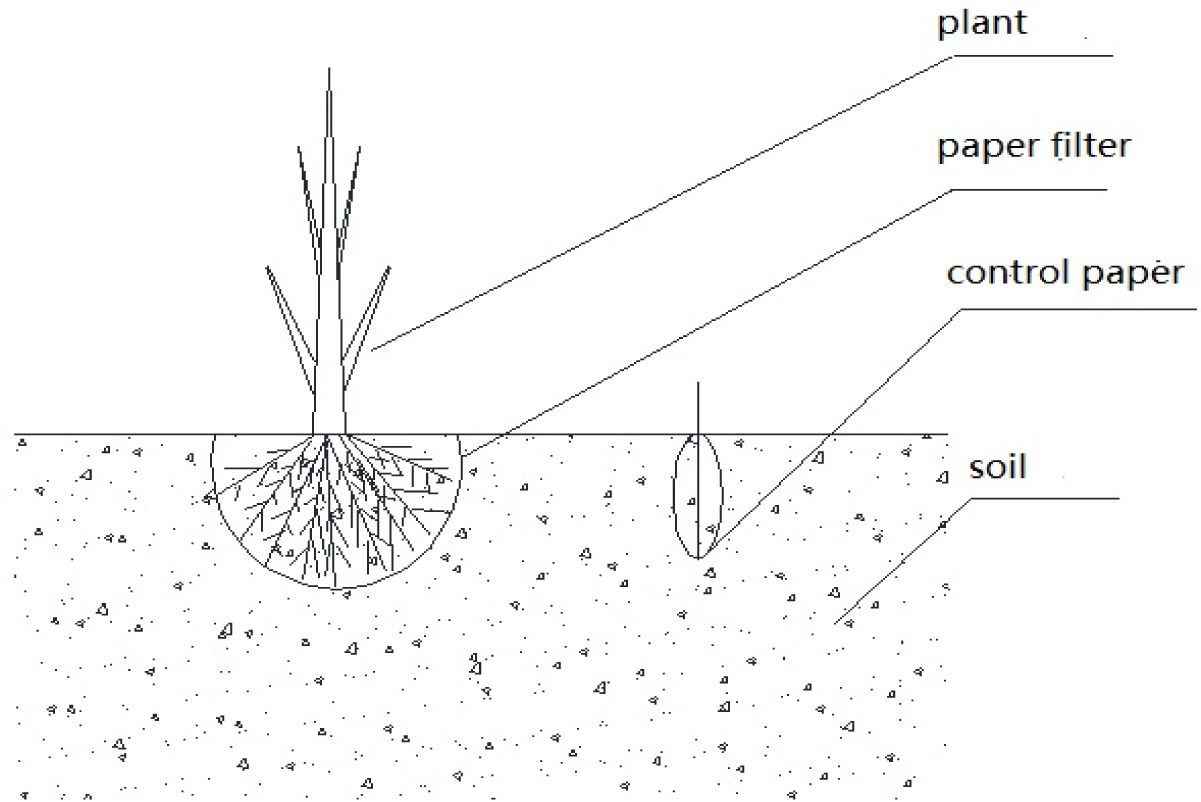
diagram of paper filter system

## Detection of quantity of objects in membrane

After several days, we will collect the filter papers to measure the materials in it. How to select the exact time will be the responsibility of the researchers. If there is not enough available information, a series of parallel experiments are suggested. For example, 3 days, 5 days, 7 days…, samples can be collected at different time point, the ‘right’ time is the time when the data is stable with the following checkpoint. We will put the two pieces of paper together and a preservative film should be put between the two pieces of the filter paper. A pair of sharp scissors are necessary to cut the two papers, which will ensure us to get the same size of filter papers. We will use gas chromatograph(or other equipment, such as HPLC, mass spectrum, depending on the materials observed) to quantify the materials in filter paper. In the end we will get N of the materials.(N=quantity(root)/quantity(soil)).

## Detect the density of interested chemicals in soil and calculating the real density circuiting the plant roots

We use the normal method to test the density in solution/soil. The density of the root will be decieded by the following equation.

As usual we perfom the experiment about allelochemicals in nutrient solution. We must convert the density in soil to the density in solution. Considering the facts that the our unique interest is the molecules, we should calculate the objective molecules number in unit volume. With this goal we should measure the density of the objective soil. The standard method is as usual^5^. With such data, we can arrive at our expected rusult, the concenration(C) of objective molecules in solution:

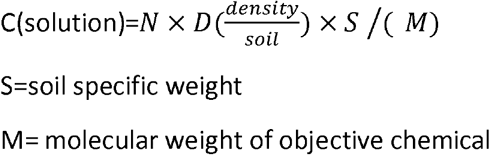

## Validation of this system

Peanut is a kind of plant we are interested in. We used huayu-22 cultivar as our research material. In the beginning we followed the routine procedure to germinate the peanut(related methodology can be found in the supplement). Afterwards we transplated it to the objective soil in which peanut is planted continuously for several years. Everyday we will water the plants to keep the soil moist. 10 days later we collected the paper filters and cut the interested part and send it to gas chromatograph. The density of soil is 1.30g/cm^3^. Benzoic acid, palmitic acid, octadecoic acid, as the most studied allelochemicals with peanut^6^, were be measured. The results are showed in table 1. With the former equation, we can either calculate the ‘real ‘density of such materials circuiting the peanut roots.

### Discussion

Filter paper is only an option in our method. Filter paper is suitable for most root ececutes in our opinion. However, for other chemicals, raw tape may be a better option. The selection should be from the suggestion of chemist.

To ensure the accuracy of measurement, we also add preservative film into the two layers of filter paper to construct a ‘sandwich’, which can avoid the interference between the two layers. Then we will cut the filter papers with the sharp scissors in the same part(figure3).

**Figure 3:**
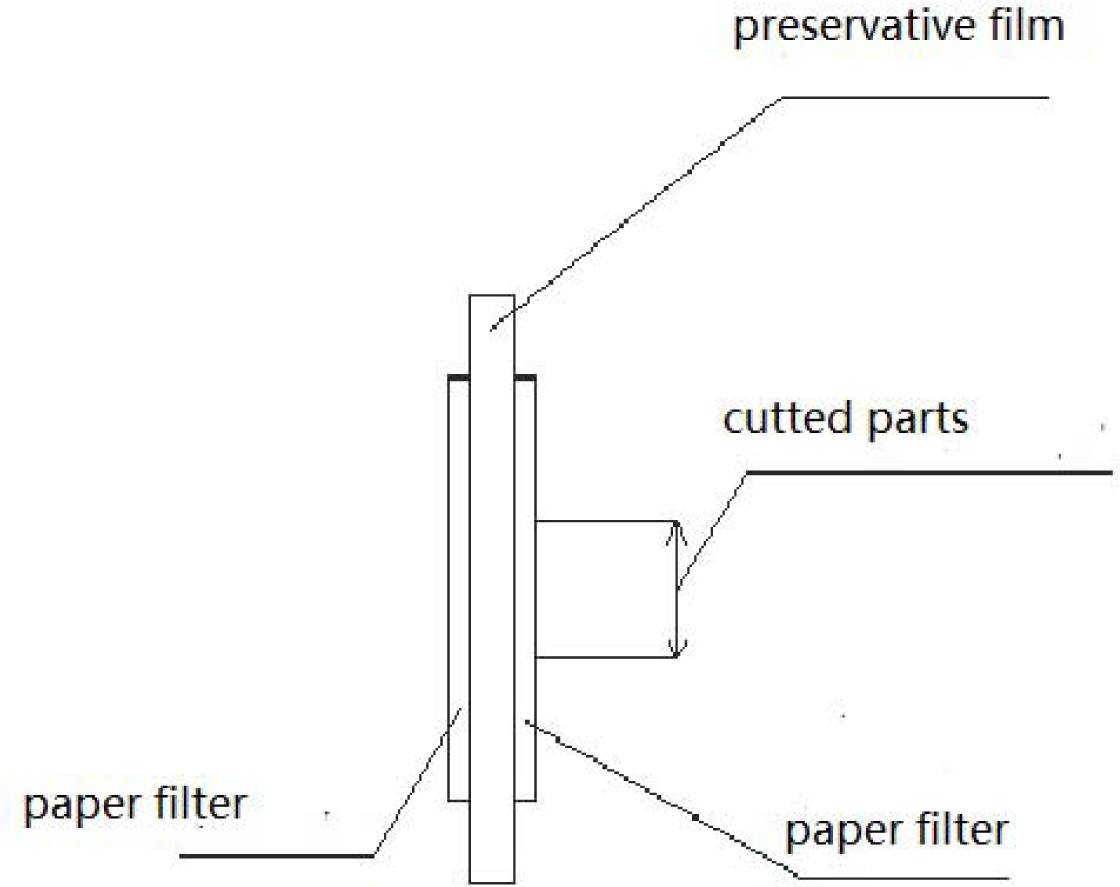
sandwich structure of paper filter sample

There exists a problem: absorbability of the filter paper will affect the calculation accuracy. For the purpose of accuracy, after the initial measurement the absorbability of filter paper(or other materials) to the objective molecules should be measured at the initial concentration. In this paper we avoid such work due to the complexity of different execute molecules and absorbing media. If adsorption coefficient A(%) is introduced into our equation, the new equation is :

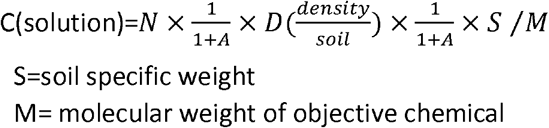

Our method has a great advantage: we can calculate the real density of root ececutes easily and fast. Moreover we can compare the density on the surface of the plant root and the concentration in the liquid media, which can help researchers decide the concentration in the liquid media when they design the experiment. Compared to the former research^7^, we can find that in spite that the concentration of chemicals in soil is low, but the density surrounding the root is very high, which will obstacle the development of peanut. Our rsults perfect explain the long-existing question: why do the low quantity of allelochemicals affect the plant development? How do the allelochemicals play their roles in crop continous obstacle?

In the future we can advocate this promising method to similar applications. For example, we also know that rhizospheric microorganism has a different constituition from the soil microoragnism. This effect also ascribes to the root executes. Many people think that plant executes will cause chemotaxis in the microorganisms. Thus this kind of chemotaxis will cause a great difference in the consitituion in microorganism. Our method will let such work can be done more precisely. The former work usually was done by a kind of rough handwork^8^

In our method, there also exists the potential to detect the precise density of object materials on the surface of different parts of the same plant roots. If we cut the sample parts from different parts of the filter paper corresponding the different parts of the root, we can detect the density of executes on the surface of different parts of root, even the microbiota.

In the former research, few work has been done about the distribution of plant root executes on the surface of root. This can be partly ascribed to the absence of a suitable method. The second reason is that the accuracy of measurement must be enough to distinguish the difference between different periods. Our method makes this kind of research possible in the future.

In spite that this method is very promising, we must develop this method to accomodate much more kinds of plants. We must acknowledge that our filter paper can not contact the roots tightly, which will lead our results lower than the factural concentration. In the meanwhile a more sensitive method than Liquid mass spectrometry is needed to apply to such detection. However, at least we can set a minimum level for liquid media when we do resear about allelopathy

