## Supplementary material for "A paper-filter system to investigate the real micro-environment circuiting plant roots": table1 and supplement method: supplements20190108.docx

online methods

peanut gemination

The peanut particles should be peeled off and we select the plump particles. Immense the particles into clean water until it geminate. Translate it to special hydroponic basin until the plant is big enough.

sample collection and gas chromatography

the filter paper(or other media) can be cut as figure 4. Then the sample should be drid in oven at 105% after 24 hr.

The following apatus and conditions are for gas chromatrophy:

Shimadzu GC-2010,column: AT-FFAP，30m*0.53mm*1.0um, temperature of Sampling port: 320℃, tempertature of column: 240℃, temperature of dector: 320℃, Sample volume: 5.0ul; column flow rate: 8.0ml/min; shunt ratio: 1:5;

control solution:

Solute 20mg obective chemical in 100ml methanol as control solution,sonicate it until soluable completely.

sample solution:

Weigh the paper filter, immense it in methanol(1.0g/20ml) for 1 hr, sonicate it until soluable completely, filt it with nylon66 filter membrane and we get the sample solution.
