## Supplementary material for "A paper-filter system to investigate the real micro-environment circuiting plant roots": table1 and supplement method: tables20181229.docx

Table1: the density of allelochemicals in soil and surrounding the peanut

|  | Quantity from paper in soil(mg/g) | Quantity from paper surrounding the roots(mg/g) | N=quantity(root)/quantity(soil) | Density in soil(mg/g) | Density surrounding root(mg/g) | Expected concentration of solution(mol/L) |
| --- | --- | --- | --- | --- | --- | --- |
| Benzoic acid | 0.001 | 0.013 | 13 | 0.001 | 0.013 | 0.0013 |
| palmitic acid | 0.003 | 0.228 | 76 | 0.007 | 0.532 | 0.0027 |
| octadecoic acid | 0.004 | 0.457 | 114 | 0.009 | 1.028 | 0.0047 |
